# Receptor dimerization enables ligand discrimination through tunable response heterogeneity

**DOI:** 10.1101/2025.01.01.630981

**Authors:** Assaf Biran, Yaron E. Antebi

## Abstract

Signaling pathways enable cells to coordinate collective behaviors by exchanging specific information. Many pathways utilize multiple ligand variants to activate the same intracellular signaling cascade, raising the question of how cells discriminate between these seemingly redundant signals. It has been shown that individual cells can discriminate between signals based on their induced level of activity, temporal dynamics or combinatorial effect. Here, we demonstrate that ligand discrimination could also emerge at the population level. Using mathematical models of ligand-receptor interactions, we examine how response heterogeneity at the population level can encode ligand identity. We introduce a local scaling metric to quantify how variation in pathway components affects the cellular response. Our results reveal that for pathways with dimeric receptors, and more significantly for heterodimeric receptors, biochemical parameters of the ligands control the resulting heterogeneity in the response of a population of cells. Furthermore, we show that the population-level heterogeneity encodes the enzymatic activity of the resulting receptor complex. This suggests a functional advantage for utilizing heterodimeric receptor complexes in pathways acting across a population of cells, such as the type I interferon pathway, which shows several of the characteristics of our model. This contrasts to juxtacrine pathways, such as Notch, that do not act at the population level and use a single component receptor. Overall, our findings highlight a novel mechanism by which receptor architecture enables cells to encode ligand-specific information through population-level heterogeneity, offering insights into immune regulation, tissue development, and synthetic biology.

**Significance Statement:** Cells often communicate using distinct molecular signals (ligands) that activate the same pathway. Yet, how cells distinguish between these signals remains unclear. This study reveals that ligand discrimination can emerge at the level of entire cell populations, not just individual cells. By analyzing mathematical models, we show that certain receptor architectures, especially those involving heterodimeric receptors, allow signals to control the heterogeneity of the response level across a population. This mechanism enables cell populations to decode specific information about their environment, impacting processes like immune responses and tissue development. Our findings provide new insights into how signals coordinate collective behaviors and suggest strategies for designing synthetic systems to precisely control biological responses.

## Introduction

Intercellular signaling pathways regulate many aspects of multicellular organisms, from development, through homeostasis, to immune responses [1–3]. These processes inherently rely on multiple cells acting together in a coordinated manner [2,3]. To achieve such coordinated behaviors robustly, cells secrete and respond to a wide set of ligands. Interestingly, many signaling pathways, such as type I interferon (IFN), bone morphogenic protein (BMP), transforming growth factor β (TGFβ), fibroblast growth factors (FGF), Notch, and others, show a peculiar feature. Multiple distinct ligands promiscuously interact with shared receptors to activate the same downstream pathway [1,4,5]. In many cases, these distinct ligands give rise to distinct biological outcomes, despite using the same intracellular mediators [6–8] (Figure 1A). Thus, a central challenge in understanding the regulatory capacity of cell-to-cell communication is determining the extent and mechanisms by which cells encode ligand identity and discriminate between seemingly equivalent ligands.

**Figure 1.**
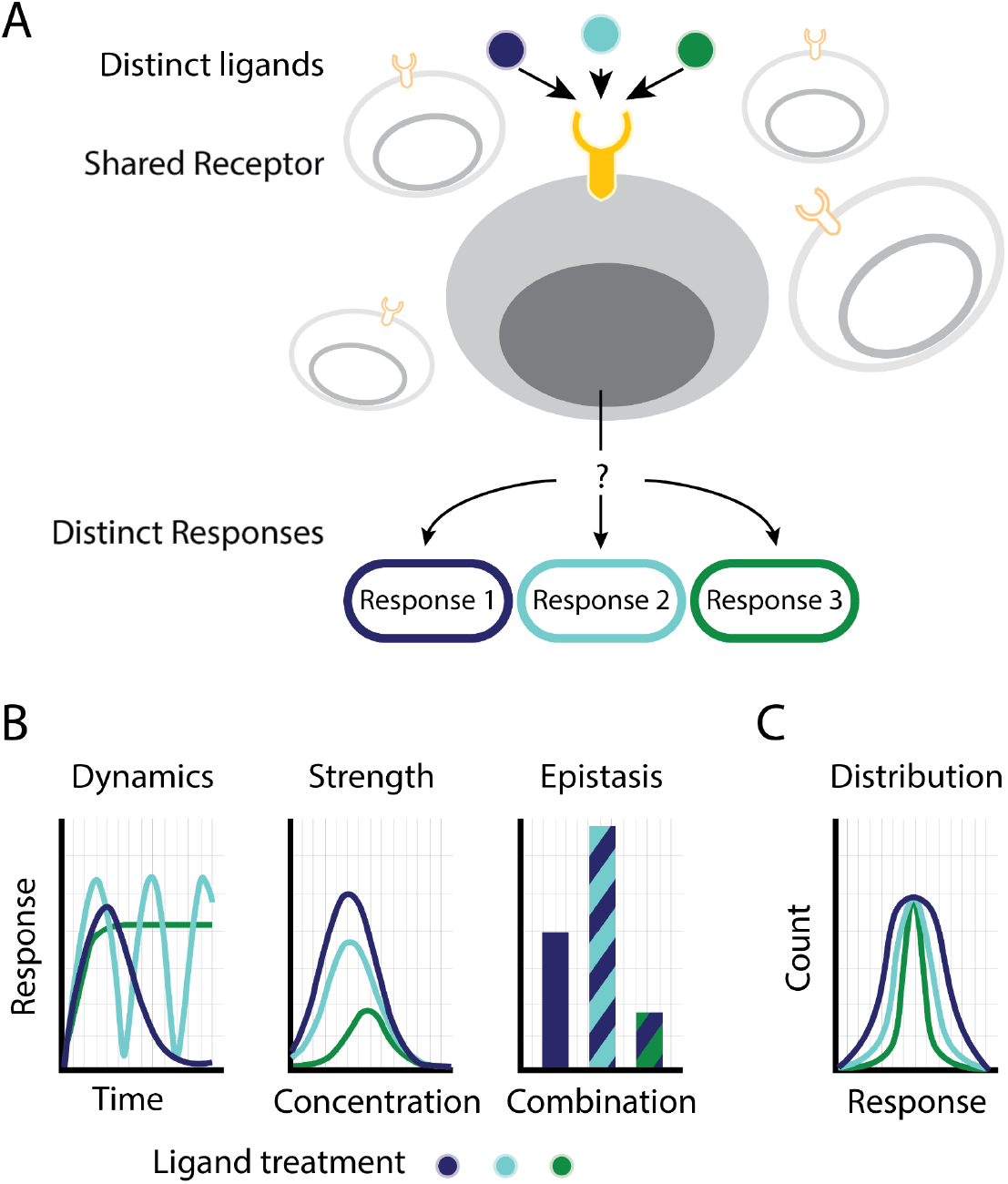
Ligand discrimination in multi-ligand pathways. (A) Signaling pathways comprise multiple ligands that bind the same receptors on the membrane of cells. This provokes the question of whether and how cells in a population discriminate between the different ligands of the same pathway. (B) Several mechanisms have been suggested for ligand discrimination on the single-cell level. In dynamic ligand discrimination, ligands elicit different response dynamics over time (left). Discrimination could also be achieved when ligands activate the pathway with distinct response strength (center). Finally, the epistatic relationship of one ligand with other ligands of the same pathway could result in a ligand-specific response (right). (C) Alternatively, ligand discrimination could occur at the level of the population. In this case, the ligand identity results in a variation of population-level properties, such as heterogeneity across the population of cells. Inducing distinct levels of variability across the tissue, ligands can result in distinct phenotypes.

Previous work has demonstrated several different mechanisms for ligand discrimination. In the Notch signaling pathway, different ligands show distinct temporal dynamics that lead to the activation of distinct downstream target genes [8] (Figure 1B, left). A similar mechanism is exhibited by the NF-KB transcription factor, which discriminates between different stimuli based on the different activation dynamics [9]. Alternatively, ligands can be distinguished through their maximal activation level (Figure 1B, middle) [6,10,11]. In the interferon pathway, ligand-specific activity strength can lead to specific expression levels of target genes [12,13] or to activation of different set of genes by activating regulatory elements with distinct affinities [14]. Lastly, it was recently shown that seemingly equivalent ligands could exhibit distinct combinatorial effects [5,15]. Such combinatorial specificity allows different ligand combinations to differentially activate specific target cells [16] (Figure 1B, right). Overall, several signal discrimination mechanisms have been identified that enable individual cells to decode the identity of the signal or signal combinations.

Many processes during development, homeostasis, and regeneration inherently control the population behavior of cells. In these cases, ligand discrimination might not arise at the single-cell level, but rather, ligands could be distinguished by the way they affect the overall population behavior. In particular, a defining trait of biological systems is the inherent stochasticity among cells. Within the same population, cells often express RNA and proteins at different levels [17–19]. This variability has been shown to have important and beneficial functional roles [17]. For example, stochasticity could result in a robust response at the population level [20,21] or allow an efficient way to regulate gene expression and differentiation [18,22]. In single-cell organisms, such as bacteria and yeast, as well as cancer cells, stochasticity in gene expression allows for bet-hedging, where a small, random subpopulation of cells express certain genes that allow them to survive extreme conditions or drug treatments [23–26]. External regulation of the stochasticity within the population has been shown to allow control over the behavior of bacteria and yeast [24,27]. Thus, ligand-dependent control of heterogeneity can have an important biological role in determining cellular responses.

A prominent example of this importance in multicellular organisms is the type I IFN antiviral pathway [3]. The type I IFN pathway is a significant component of the innate immune response. it helps protect against infections and has an immunomodulatory role in cancers [3,12]. Importantly, excess activation of the type I IFN pathway was found to be involved in the initiation and sustainment of autoimmune diseases [28,29]. As such, it is critical for the number of responding cells to be tightly controlled [28]. It was further found that cell-cell variability in this pathway originates from heterogeneity in the amount of receptor expressed [12–14]. This suggests a new type of ligand-specific effect in which cells can be activated at the same average level but with distinct levels of heterogeneity across the population in a ligand-dependent manner (Figure 1C). In this way, different ligands could control the variability of the response within the population, leading to distinct systematic behaviors.

Here we use mathematical models, motivated by the type I IFN pathway, to analyze the capacity of signaling pathways to exhibit ligand-dependent heterogeneity. We formulate and analytically solve models describing several receptor architectures. For each architecture, we analyze the response heterogeneity and determine to what extent ligand identity can attenuate this heterogeneity. We start with models of single subunit receptors and find a robust heterogeneity that cannot be controlled by extracellular ligands. Continuing to study a more complex model of receptors acting through ligand-induced dimerization, we find that heterogeneity depends on the total amount of full complexes that are formed. In this case, ligand parameters can affect and tune the relative heterogeneity. Moreover, we distinguish between homo- and heterodimerizing receptors. Models with homodimerizing receptors show limited sensitivity to ligand characteristics, giving rise to a limited ligand discrimination capacity. Models with heterodimerizing receptors, however, allow for greater control over the response heterogeneity in a ligand-dependent manner. As such, they provide a mechanism for a flexible ligand discrimination capacity. Taken together, our results reveal the impact of ligand identity on the distribution of cell response to stimuli for different receptor architectures. In pathways with multi-component receptor complexes, such as type I IFN and other immunological and developmental pathways, this suggests that specific ligand variants can result in distinct overall heterogeneity in the response that can lead to ligand-specific functional phenotype.

## Results

### Signaling pathways with monomeric receptors give rise to response heterogeneity that is independent of ligand identity

We start by considering the basic architecture of a signaling pathway, which consists of a single-unit receptor that binds an extracellular ligand. The formation of a ligand-receptor complex then leads to the induction of a downstream intracellular response. Such architectures are utilized in several biological pathways, such as the Notch signaling pathway [8], and certain G-protein coupled receptor systems, such as the CCR1 chemokine receptor and its ligands [30]. Such systems can be described using a minimal mathematical model, where a ligand, denoted by *L*, binds to a receptor, *A*, to form a fully active signaling complex, *F* (Figure 2A). Once a complex is formed, it induces intracellular changes leading to the transcription of target genes. The behavior of this model is governed by four parameters. The total amount of receptors on the cell surface, A^0^, is inherent to the cells and independent of the extracellular environment. In contrast, the concentration of the ligand in the environment, *C*^*0*^, the affinity of the ligand to the receptor, *K*_*L*_, and the rate by which the complex activates the downstream pathway, *e*_*L*_, are properties of the specific ligand used and can be changed accordingly. At steady-state, the model can be specified by the set of ligand-receptor binding-unbinding equations, together with an additional equation for mass conservation of the total receptors (supplementary text). Here, we assume that the ligand is in excess, and as such, the binding of ligands to the receptor does not affect their overall concentration in the environment. Solving these equations at steady-state gives rise to the Michaelis-Menten dependence of response on ligand concentration (Figure S1A) [6,10].

**Figure 2.**
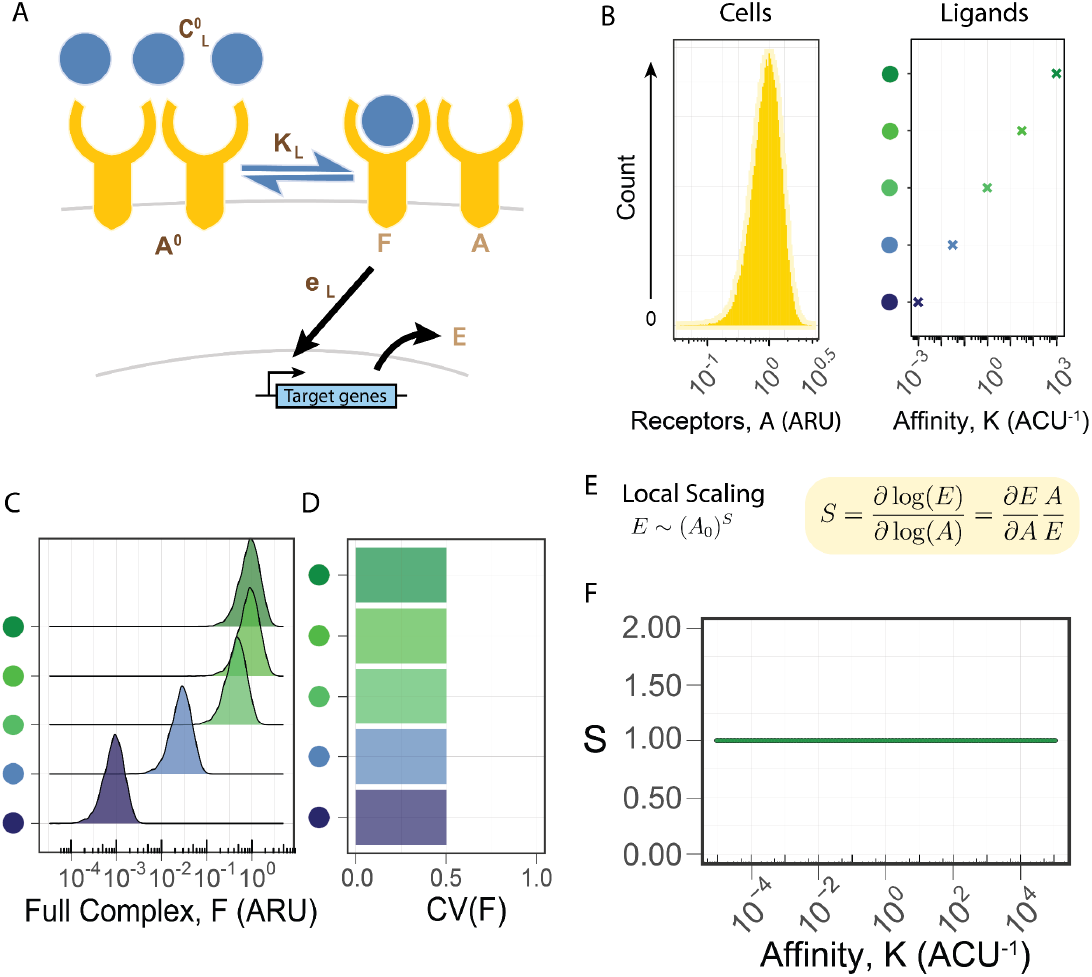
Single-unit receptor architecture does not allow for ligand-dependent heterogeneity. (A) A minimal model for a pathway with a single-unit receptor. Receptors (A) and ligands (C) bind to form full complexes (F) and activate target genes (E). The four model parameters (total receptors, A^0^; ligand concentration, C^0^; affinity, K; and enzymatic efficiency, e) are shown in dark brown, and the three variables are shown in light brown. Parameters that depend on the specific ligand identity are denoted with a subscript L. (B) We consider the response of a population of 10,000 cells with a Gamma distribution of receptors with a mean of one (ARU = Arbitrary Receptor Units) and standard deviation of 0.5 (left) to five ligands with a concentration of one (ACU = Arbitrary Concentration Units) and different affinity values (right). (C, D) The response distribution (C) and the coefficient of variation (D) are plotted for each ligand. (E) The local scaling, defined as the logarithmic derivative, reflects the relative change in the response to relative changes in receptors. (F) The scaling with the single-unit receptor model is constant and independent of the ligand parameter.

Using the analytical solution, we can determine the response heterogeneity and its dependence on ligand identity through its chemical properties. In our analysis we quantify heterogeneity by the coefficient of variation (CV) to normalize the general dependence of standard deviation on rescaling of the distribution. To analyze the heterogeneity, we assume that the media is well mixed, and thus all cells receive the same ligand concentration. We focus on response variability derived from noise in receptor expression levels. This has been shown to account for most of the variability in the IFN type I pathway [31]. We considered a population of 100,000 cells and simulated variable receptor expression levels distributed with a gamma distribution (Figure 2B, left). We examine five different ligands at the same concentration but with different affinities (Figure 2B, right). While the mean amount of formed complexes is dependent on the ligand parameters (Figure 2C), the heterogeneity is robust across all ligands, regardless of their affinity to the receptor (Figure 2D). These examples suggest that in this model, the response heterogeneity is independent of any biochemical parameter and cannot be adjusted.

We next aimed to determine the robustness of response heterogeneity more systematically and for general distributions of receptors. Using simulations to compute distinct distributions of receptors with distinct levels of variability is computationally prohibitive [21]. Instead, we pursued a more basic metric for analyzing heterogeneity that is independent of the specific shape of the overall distribution. We consider the local scaling, S, between the response and the amount of receptors in a specific cell (Figure 2E, S1B, supplementary text). This scaling is defined to be the relative change in the response (E) for a small relative change in the receptor amount (A_0_) and represents the power law dependence of the response on the amount of receptors (E∼A_0_^S^). A system with a linear dependence, S=1, will have a similar variability in the response as in the receptors. For a sublinear dependence, S<1, the CV of the response is reduced compared to the CV of the receptors, while a superlinear dependence, S>1, results in increased response heterogeneity. We are interested in determining whether S depends on ligand parameters, in which case the response variability will depend on the identity of the ligand. Thus, this local scaling metric allows us to analytically determine the capacity of ligand identity to determine population heterogeneity in the response.

Computing the local scaling metric for the basic model of a single unit receptor, we find that the scaling is identically equal to one, independently of any biochemical parameter (Figure 2F). This reflects the linear dependence of the response on the receptors (supplementary text) as each receptor binds independently to the ligand, and its contribution to the total complex amount is independent and additive. In this case, the scaling does not depend on the parameters and the response heterogeneity reflects only the heterogeneity of the receptors, in agreement with our simulation-based result (Figure 2C, D). We thus find that in a single-unit receptor system, the population distribution does not provide a way to discriminate between ligands.

### In pathways with dimeric receptors, ligand properties can affect the response heterogeneity

Pathways with a single unit receptor are not generally common in mammalian systems. Rather, in most mammalian signaling pathways, receptors are formed from a bound complex with multiple receptor subunits. In some cases, e.g., the FGF pathway and other receptor tyrosine kinases, a ligand binds to two equivalent receptor subunits, which form a full signaling complex and initiate the intracellular response [32–34]. To study this more complex architecture of homodimeric receptors, we extended our model to describe a two-step formation of the signaling complex (supplementary text). Briefly, a ligand denoted by *L* first binds to one receptor subunit, *A*, to form a partial complex *P*_*L*_ with an affinity of *K*^*P*^_*L*_. Once *P*_*L*_ is formed, it can bind a second identical receptor subunit, *A*, to form the full, tetrameric complex *F*_*L*_, with an affinity *K*^*F*^_*L*_. The full complex, *F*_*L*_, can then induce the expression of target genes, *E*, at a rate *e*_*L*_ (Figure 3A). In this model, the ligand-dependent parameters are the concentration of the ligand, *C*_*L*_, its affinity to the receptor subunit, *K*^*P*^_*L*_, the affinity of the dimeric complex, *K*^*F*^_*L*_, and the activation rate of the full complex, *e*_L_. Together, these four parameters define a specific signaling environment for the cells. The configuration of the cells is defined by the level of total receptor subunits, *A*^*0*^, which does not depend on the ligand identity. As before, we assume that ligands are in excess and, therefore, their concentration is not reduced upon binding of the receptors. This extended version of the model allows us to analyze the more complex case of dimeric receptors.

**Figure 3.**
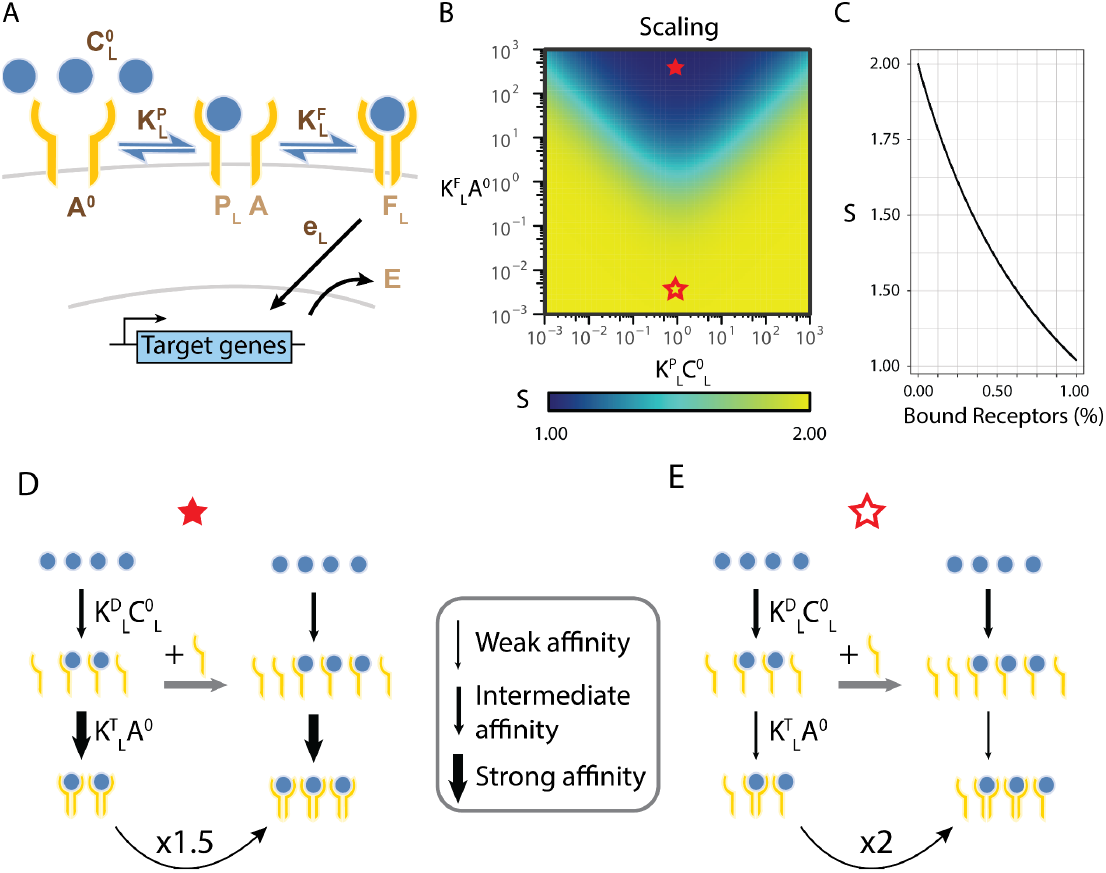
Homodimeric receptor architecture allows for ligand-dependent scaling. (A) A minimal model for a pathway with a homodimeric receptor. Receptor subunits and ligands bind to form partial and full complexes and activate target genes. The five model parameters are shown in dark brown, and the four variables are shown in light brown. Parameters that depend on the specific ligand identity are denoted with a subscript L. (B) The model’s scaling (*S*) to changes in the receptor subunit *A*^*0*^ is plotted as model parameters vary. The full and hollow red stars indicate the parameter regimes discussed in D and E, respectively. (C) The dependence between the scaling and the fraction of bound receptors (2*F*_*L*_/*A*^*0*^) is plotted. (D, E) The effect of receptor addition on the response depends on the model’s parameters. Arrow thickness indicates the relative strength of the parameters. (D) When *K*^*F*^_*L*_*A*^*0*^*>>1* and *K*^*P*^_*L*_*C*^*0*^_*L*_≈*1* (full star in B), all free receptor subunits *A* bind to any free partial ligand-receptor complex *P*_*L*_ so that the amount of *P*_*L*_ *is* the limiting factor in complex formation, and *F*_*L*_ is linearly dependent on *A*^*0*^. (E) When *K*^*F*^_*L*_*A*^*0*^*<<1* and *K*^*P*^_*L*_*C*^*0*^_*L*_≈*1* (hollow star in B), most *P*_*L*_ will be free of receptors. In this case, *F*_*L*_ will be proportional to the product of *A* and *P*_*L*_, and as *P*_*L*_’s amount is proportional to that of *A, F*_*L*_ scales quadratically with *A*^*0*^.

We next used the model to determine the scaling of the response to changes in receptor levels. We start by focusing on binding-unbinding parameters, keeping the rate of downstream activation fixed, *e*_*L*_ = 1, for all ligands. Using dimensional analysis (supplementary text), we find that the behavior of the system depends only on the dimensionless quantities *K*_*L*_^*P*^ *C*_*L*_ and K^F^_*L*_ *A*. This is consistent with previous studies of this model, which have shown that the ligands’ concentration can be compensated for by its affinity to the free subunit, *K*^*P*^_*L*_, and that the amount of receptors can be compensated for by the dimeric complex’ affinity to the free subunit, *K*^*F*^_*L*_ [35,36]. Solving the response in steady-state gives rise to a non-monotonic dependence on ligand concentration (Figure S2A). Using the expression for the response, we analytically solved for the local scaling (Figure 3B, C, supplementary text). In contrast to the single-unit receptor model, we find that the scaling depends on ligand parameters but has a limited range, showing a fold-change of two.

The model can further determine the molecular mechanism allowing controlled heterogeneity in this context and gaining an intuition for its limited range. To this end, we looked more closely at the different parameter regimes of the model. We focus on two ligand parameter regimes exhibiting distinct scalings, with one ligand having the lowest possible scaling (S=1, linear dependence) and the other the highest (S=2, quadratic dependence) (Figure 3B, filled and empty star). The first regime (marked by a filled star) is characterized by high trimeric affinity (*K*^*F*^_*L*_*A*^*0*^≫*1)* and a concentration of ligands around the EC50 for the formation of dimers *K*^*P*^_*L*_*C*_*L*_≈*1*. In this regime, the equation for the dependence of the activity on the receptors simplifies and becomes linear (supplementary text). Intuitively, if we start with free receptors, the first binding reaction is at its EC50 and results in half of the receptor forming dimers while the other half of them will remain free. Since the formation of the full trimeric complex is saturated (*K*^*F*^_*L*_*A*^*0*^≫*1*), all dimers will then bind with the available receptors.

Overall, most receptor subunits will be paired up efficiently to form full complexes, resulting in a linear dependence of the total amount of complexes, *F*_*L*_, on the amount of receptor subunits, *A*^*0*^ (Figure 3D).

In the second regime (empty star), on the other hand, the full complex affinity is low (*K*^*F*^_*L*_ *A*^*0*^*<<1*). In this case, the second step reaction, forming the full complex, is slow and far from saturation, resulting in many available free dimers and free receptors. In this regime, forming a few full complexes does not significantly change the amount of receptors and dimers. In other words, this regime can be analyzed as a binding reaction with no depletion of the components. In such conditions, the amount of full complexes will be proportional to the product of the amount of receptors with the amount of dimers. Considering the EC50 concentration for dimer formation, the amount of dimers is proportional to the amount of receptors. Thus, the overall amount of full complexes, *F*, scales quadratically with the amount of receptors, A^0^ (Figure 3E). This can also be shown analytically to hold, even for a more general range of parameters (see supplementary text). We see that the dependence of the amount of full complexes on the amount of receptors shifts from linear in the first regime to quadratic in the second. Accordingly, the local scaling of the full complex, with respect to receptor levels, varies from one to two, reflecting a corresponding change in the heterogeneity.

### Response heterogeneity encodes the relative effective activity parameter of the ligand

We next studied what ligand properties are important for controlling the heterogeneity. We found that while the scaling has a complex dependence on the parameters, it can be written as a simple function of the full tetrameric complex, F_L_, with no additional dependence on parameters (Figure 3C, supplementary text). When more complexes are formed, the system becomes less sensitive to changes in receptors, and the scaling is reduced. We emphasize that this inverse dependence emerges from the architecture of the pathway, as it does not occur for the simpler, monomeric receptor architecture.

The direct dependence of the scaling on F_L_ has an important implication. While the overall response depends both on the amount of complexes as well as the activity rate, *e*, the scaling does not depend on the activity rate (supplementary text). Hence, if cells are exposed to ligands with distinct activity parameters, then for the same overall mean response, the population will necessarily exhibit distinct complexes amount which would mean a difference in the response variability. In this way, a dimeric system encodes the activity rate of ligands into variability in the response, and thus the population response can differ between ligands, even when the mean response is the same.

This population-level effect can be demonstrated by using the model to simulate the response of a population of cells under ligands with distinct parameters. To this end, we simulated a population of 100,000 cells with gamma-distributed expression of receptor *A*, as before. We analyzed the response’s distribution under the induction of five different ligands with different parameters (Figure 4A). For each ligand, we chose the parameters such that the mean response is identical while the activity rate differs. Calculating the CV, as before, we find that the different ligands induce a different level of heterogeneity in the response, even though they have the same response mean (Figure 4B, C). As we observed for the scaling metric, the ratio between the CV across different ligands is limited to twofold. Furthermore, we see that the standard deviation of the response can be closely approximated by the product of the receptor standard deviation by response scaling (Figure S2B). Thus, ligand-dependent variability can be seen for simulated distributions of cells.

**Figure 4.**
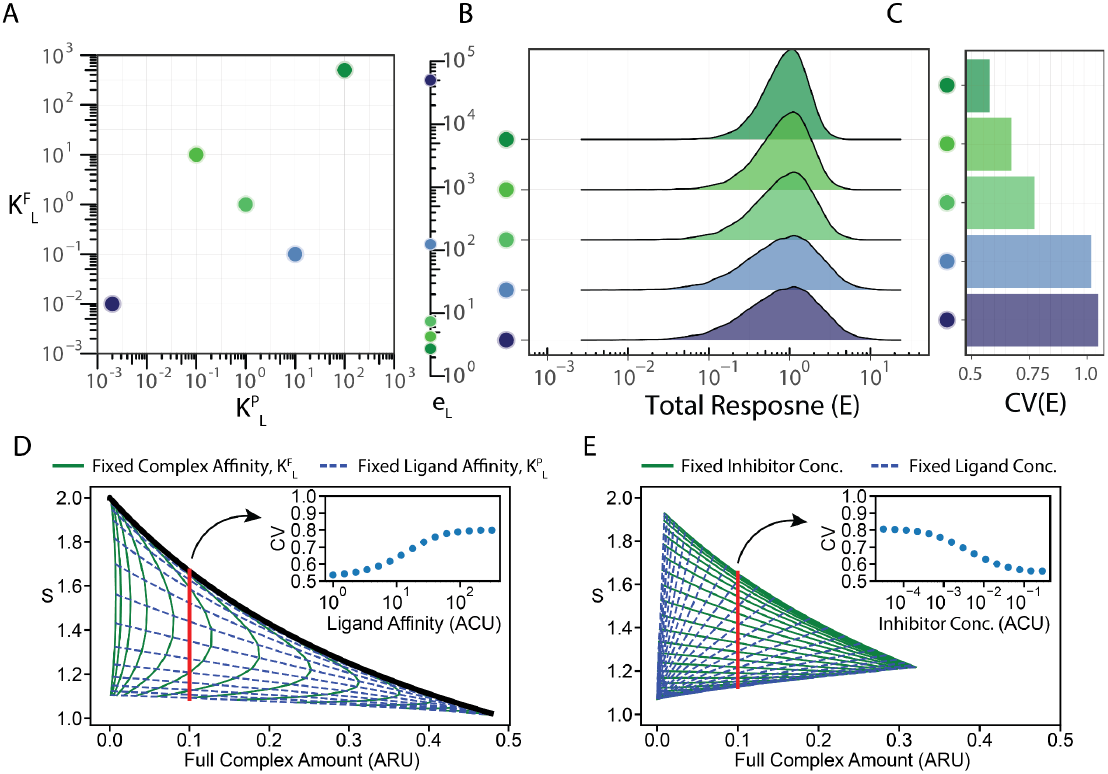
Ligand efficiency is encoded in the heterogeneity of the response. (A) We consider five ligands with different affinities (*K*^*F*^_*L*_, *K*^*P*^_*L*_) and activation rates (*e*_*L*_). Different parameters were chosen with a fixed mean response of one. (B, C) We simulated the response of a population of 100,000 cells to the five ligands. Cells were assumed to express receptors with a Gamma distribution of unit mean and 0.5 standard deviations. The response distribution (B) and the coefficient of (C) are plotted for each ligand. (D)The scaling was simulated for ligands with different chemical parameters in the presence of a zero-activity inhibitor. A green full line represents some fixed complex affinity, K^F^_L_, with varying ligand affinity, K^P^_L_. A blue dashed line represents some fixed ligand affinity with varying complex affinity. The black line represents the dependence without an inhibitor (cf Figure 3C). A set of affinity parameters resulting in a fixed response level (red line) was selected, and the CV of the response was computed for a population of cells as before (inset). (E) The scaling was simulated at varying ligand and inhibitor concentrations. A green full line represents some fixed inhibitor concentration with varying ligand concentrations. A blue dashed line represents some fixed ligand concentrations with fixed inhibitor concentrations. A set of relative concentrations resulting in a fixed response level (red line) was selected, and the CV of the response was computed for a population of cells as before (inset). ARU = Arbitrary Receptor Units, ACU = Arbitrary Concentration Units.

An extreme example of ligands with different activity rates is the case of competitive inhibitors. These inhibitors can bind the receptor but do not activate the downstream pathway. In our model, this is the case where e_L_=0. Since for this case, the pathway remains completely off, we decided to analyze the case of a combination of different ligands with an inhibitor. As this two-ligand model does not lend itself to an analytic solution, we simulated the system using the EQTK toolbox [37,38]. We varied the two affinity parameters for the activating ligand while keeping the inhibitor fixed. We find that the presence of an inhibitor can break the direct dependence between S and F_L_ (Figure 4D). Distinct ligands show distinct scaling properties, even with the same efficiency. When simulating the response of a population of cells, we can see that, indeed the CV depends on the ligand affinity (Figure 4D, inset). Finally, we note that changing the concentration of the inhibitor and the signal also results in different scalings for the same number of complexes (Figure 4E). In this way, competitive inhibitors provide a mechanism to tune the heterogeneity of a response. If we vary the amount of inhibitor and signal such that the number of complexes stays constant, we see that the heterogeneity across the population is reduced as the amount of inhibitor increases (Figure 4E, inset). Our results demonstrate a new role for competitive inhibitors in shaping the heterogeneity of the response to signals.

### Heteromeric receptor systems allow general tuning of response heterogeneity

Dimerization of receptors does not always occur with two identical receptor subunits. In fact, many pathways, such as the BMP, TGFβ, and type I IFN pathways, utilize two types of receptor subunits to form an inherently heterodimeric complex [4,5,39]. We tested whether such architectures could lead to a more substantial control over the response heterogeneity compared with the two-fold limit arising in the homodimeric model. To explore this case, we added a second receptor subunit *B*, into our model (Figure 5A). Briefly, a ligand, denoted by *L*, first binds to a specific receptor subunit, denoted by *A*, to form a partial complex, *P*_*L*_, with an affinity of *K*^*P*^_*L*_. Only once *P*_*L*_ is formed it then binds the second subunit, denoted by *B*, to form the full tetrameric complex, *F*_*L*_, with an affinity *K*^*F*^_*L*_. As with the previous models, once the full complex *F*_*L*_ is formed, it will induce the expression of target genes, *E*_*L*_, in a ligand-dependent activity rate, *e*_*L*_. This model represents the current knowledge for the type I IFN, and TGFβ pathways, which canonically have only one pair of receptors, and the affinity hierarchy results in a sequential complex formation [3–5,39]. Importantly, while it is sometimes possible for a ligand to bind either of the two subunits first *in-vitro* [40,41], when testing *in-vivo*, the ligand binds preferentially to one subunit, resulting in an effective sequential assembly of the full receptor [4]. Allowing the ligand to bind either receptor subunit first does not change the main conclusions (supplementary text).

**Figure 5.**
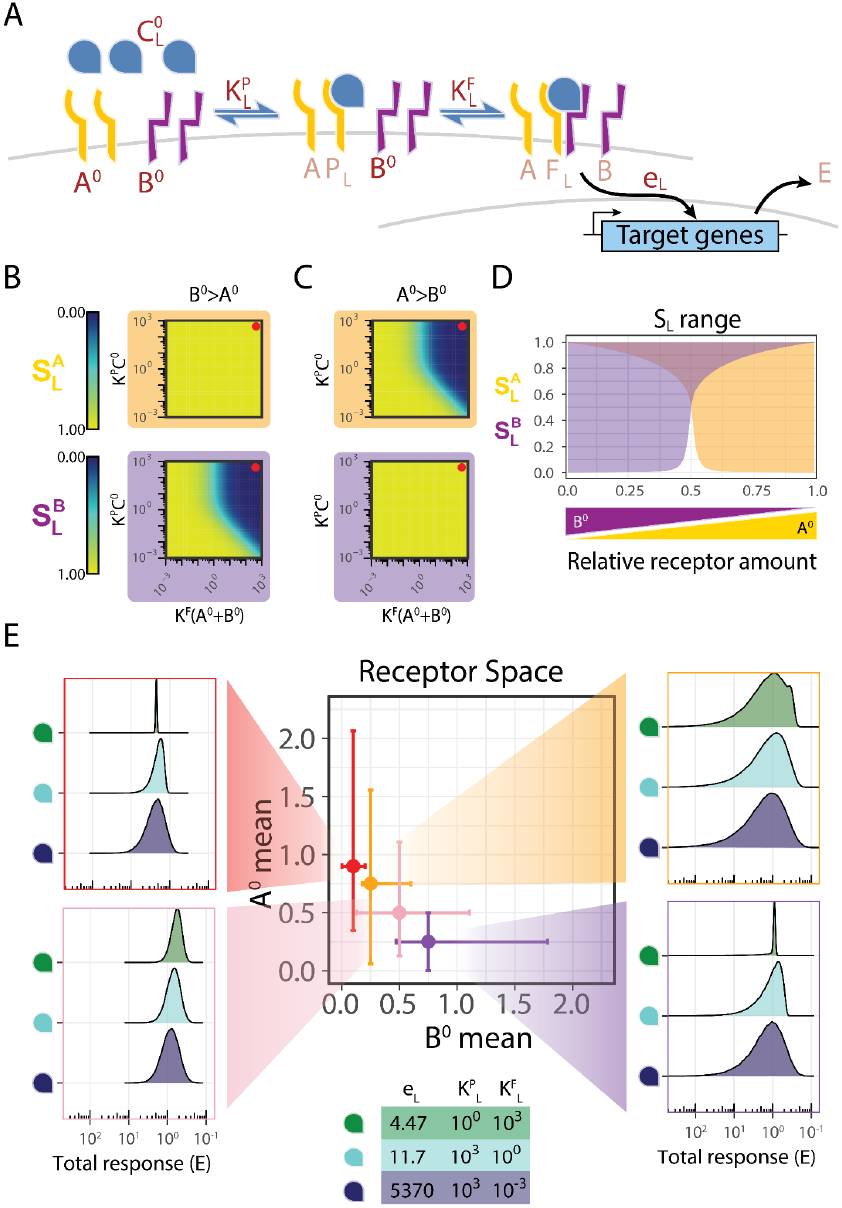
Heterodimeric receptor architecture allows for arbitrary heterogeneity-based ligand discrimination. (A) A minimal model for a pathway with a sequentially formed heterodimeric receptor. Receptor subunits and ligands bind sequentially to form partial and full complexes and activate target genes. The six model parameters are shown in dark brown, and the five variables are shown in light brown. Parameters that depend on the specific ligand identity are denoted with a subscript L. (B, C) The scaling of the model in the levels of each receptor subunits were computed across parameters for the case of *A*^*0*^<*B*^*0*^ (B) or *A*^*0*^>*B*^*0*^ (C). The scaling with the A subunit (*S*^*A*^_*L*_) is plotted with an orange background, while the scaling with the B subunit (*S*^*B*^_*L*_) is plotted with a purple background. The red circles indicate parameter regimes discussed in Figure S4A, B. (D) The ranges of the scaling levels (*S*^*A*^_*L*_ in orange and *S*^*B*^_*L*_ in purple) achievable across ligand parameters are plotted for different relative levels of the receptor subunits. (E) We consider four populations (red, yellow, pink, purple) of 100,000 cells with a Gamma distribution of receptor subunits with different means and standard deviations. The means and standard deviations for *A*^*0*^ are [0.25, 0.5, 0.75, 0.9] and [0.5, 0.25, 0.75, 0.5] respectively, and for *B*^*0*^ are [0.75, 0.5, 0.25, 0.1] and [0.1, 0.25, 0.001, 0.01]. We simulated the response of each population to three ligands with different affinity values (bottom). For each ligand, the activity rate was set so that the mean response would be one.

The model is described by six biochemical parameters. The ligand-dependent parameters are the concentration of the ligand, *C*_*L*_, its affinity to the free subunit *A, K*^*P*^_*L*_, the affinity of the dimeric complex *P*_*L*_ to *B, K*^*F*^_*L*,_ and the activity rate of the resulting complex, *e*_*L*_. The set of these four parameters together defines a specific signaling environment for the cells. In addition, the configuration of the cells is defined by the level of the receptors. Here, we will parameterize this configuration using the total receptor subunits and their ratio, A^0^/B^0^. As before, we assume that ligands are in excess so that their concentration is not significantly affected by binding to the receptors. We start by considering the case *e*_*L*_=1 for all ligands and solving the model in a steady state. Using dimensional analysis, we find that the system’s response depends on three-parameter combinations (Figure S3A, supplementary text).

In order to determine the capacity of ligand identity to determine response heterogeneity, we computed the scaling for this model by considering changes in each of the two receptor subunits, A and B, independently (Figure S3A-D, supplementary text). We find that the scaling can vary with the ligand’s biochemical parameters, indicating a tunable heterogeneity in the response, as for the homodimeric model. However, we found that in the regime where there is an imbalance between the expression levels of the two receptors, the scaling can have arbitrarily low values (Figure 5B-D, Figure S3E), in contrast to the two-fold limit found in these results, the homodimeric model. Specifically, the scaling related to the more abundant receptor shows stronger dependence on the ligand parameters, while the scaling related to the less abundant receptor is constrained to be around one. These results hold independently of the total number of subunits on the cell surface and on the values of the activity rate parameters, *e*_*L*_ (Figure S3F-G, supplementary text). As in the case of homodimeric receptor architectures, we find the total S, as well as SA and SB, are functionally dependent directly on F_L_ (Figure S3E). Overall, we find that an architecture of heterodimeric receptors enables ligands to tune significantly the scaling parameters.

Using the model, we can get intuition about the molecular mechanism allowing two distinct receptors to have more flexibility in controlling the response variability. We can consider a parameter regime where both dimeric and trimeric affinities are low (*K*^*F*^_*L*_*(A*^*0*^+*B*^*0*^*)<<1, K*^*P*^_*L*_*C*_*L*_*<<1*), and thus, both complexes *P*_*L*_ and *F*_*L*_ are sub-saturated. In this regime, the number of complexes depends linearly on the level of each receptor subunit. We note that this is analogous to the regime with quadratic dependence in the previous model. To see how the scaling changes, we compare this to a second regime where both affinities are high (*K*^*F*^_*L*_*(A*^*0*^+*B*^*0*^*)>>1, K*^*P*^_*L*_*C*_*L*_*>>1*), and thus all the binding reactions are saturated (Figure 5B, C, red circles). In this case, all receptors form complexes as long as there are enough free receptor subunits, and the scaling with respect to a particular subunit depends on its relative abundance. In the case where *B* is the more abundant (Figure S4A), both *A* and *P*_*L*_ are saturated. Thus, any addition of ligand *L* will directly bind to *A* to form *P*_*L*_, which will bind to *B* to form the full complex. Accordingly, the response will be sensitive to changes in the amount of *A* (Figure S4A, right) as the limiting subunit. In contrast, as there are many more *B* subunits than *A*, most of them would remain free, not affecting the amount of *F*_*L*_ and making the response insensitive to changes in B (Figure S4A, left). When *A* is more abundant (Figure S4B), *P*_*L*_ dimers will form, but most of them will not be bound by B. Accordingly, while changes to *A* would affect the amount of *P*_*L*_, it wouldn’t affect the amount of *F*_*L*_, and the response would be insensitive to these changes (Figure S4B, right). However, any addition of *B* would directly bind to *P*_*L*_, increasing the amount of *F*_*L*_ (Figure S4B, left). As such, the response in this regime is sensitive to changes in *B*. This regime extends the regime with linear dependence in the homodimeric model. For intermediate parameters, the dependence of the response on the abundant subunit will be sublinear. Overall, the distinct identity of the two subunits results in a sublinear dependence of the response on each receptor, which provides for the large fold-change range of the scaling metric.

### Response heterogeneity shows a larger tunability range when the expression of limiting receptors varies across cells

In order to determine how the extended range of the scaling metric affects the overall variability in the response of cell populations, we simulate the response across a cell population, as described before. However, as opposed to the previous receptor-ligand architectures, where receptors comprise a single protein variant, here there are two receptor subunits. Thus, we performed several simulations of 100,000 cells for distinct mean expression levels for the subunit receptors *A* and *B* and for distinct levels of expression variability (Figure 5E). For each population of cells, we analyzed the response distribution under the induction of three different ligands with different parameters. As with the previous model, in order to focus on the heterogeneity of the response, we adjusted the parameters such that the mean of the response is identical across ligands. We calculated the coefficient of variation for each cell population and found that the response heterogeneity varies within a large range across the distinct simulated populations. This effect is observed when there is a large variation in the expression of the two receptors and when the more abundant receptor shows larger heterogeneity across the population of cells. For these parameters, we find more than an order of magnitude difference in the response heterogeneity between ligands. When the receptor expression levels are outside this regime, the difference in heterogeneity induced by the ligands converges back to a twofold difference. Indeed, our results for the specificity show a stronger dependence on ligand parameters when the ratio of the abundances of the two receptor subunits is away from one. More specifically, the scaling relative to the more abundant receptor is tunable, and variability in that receptor will contribute to a ligand-dependent variation. In contrast, the scaling with respect to the less abundant receptor is close to 1, and large variability in that receptor will give a ligand-independent contribution to the response variability. Thus, we see that the response heterogeneity in the simulation reveals a behavior consistent with the analytical behavior of the scaling, allowing for large differences in the heterogeneity generated by different ligands.

## Discussion

Variability in biological systems has been shown to play a significant and functional role [17,18,20,24]. Thus, externally regulating the heterogeneity within a population of cells can have an important effect on biological processes [42,43]. While extracellular signals are known to control the level of intracellular activity through the signaling pathways, effects that could control the heterogeneity of the response across cells have not been studied extensively. Here we show that for specific architectures of receptor-ligand interactions, the response heterogeneity can be directly controlled by the specific ligand, with different ligands acting through the same pathway resulting in the same mean activity but with different degrees of heterogeneity.

We employed mathematical modeling to quantify the dependence of the response heterogeneity on ligand-dependent parameters. A new local scaling metric that quantifies response susceptibility to changes in pathway components enabled us to perform a local analysis of the variability in the response without assuming any specific form of the overall heterogeneity in the component. We found that pathways based on single unit receptors provide a robust and parameter-independent heterogeneity in steady-state. Thus, any two ligands, at concentrations that activate the pathway to the same mean level, result in the same population level variability as well (Figure 6A). Extending our analysis to additional receptor architectures, we found that signaling pathways initiated by receptor dimerization can result in ligand-dependent heterogeneity. In these cases, two ligands can produce the same mean pathway activity while having distinct population heterogeneity. In particular, systems with homodimeric receptors allow for only limited differences in the heterogeneity of up to a two-fold difference in the population standard deviation (Figure 6B). However, systems with heterodimeric receptors allow for a larger fold range of heterogeneity levels when one of the receptor subunits is highly expressed and more variable (Figure 6C). Significantly, we find that in these cases, the response heterogeneity can be attenuated in a ligand-dependent manner providing a novel approach for ligand discrimination at the population level rather than by any individual cells.

**Figure 6.**
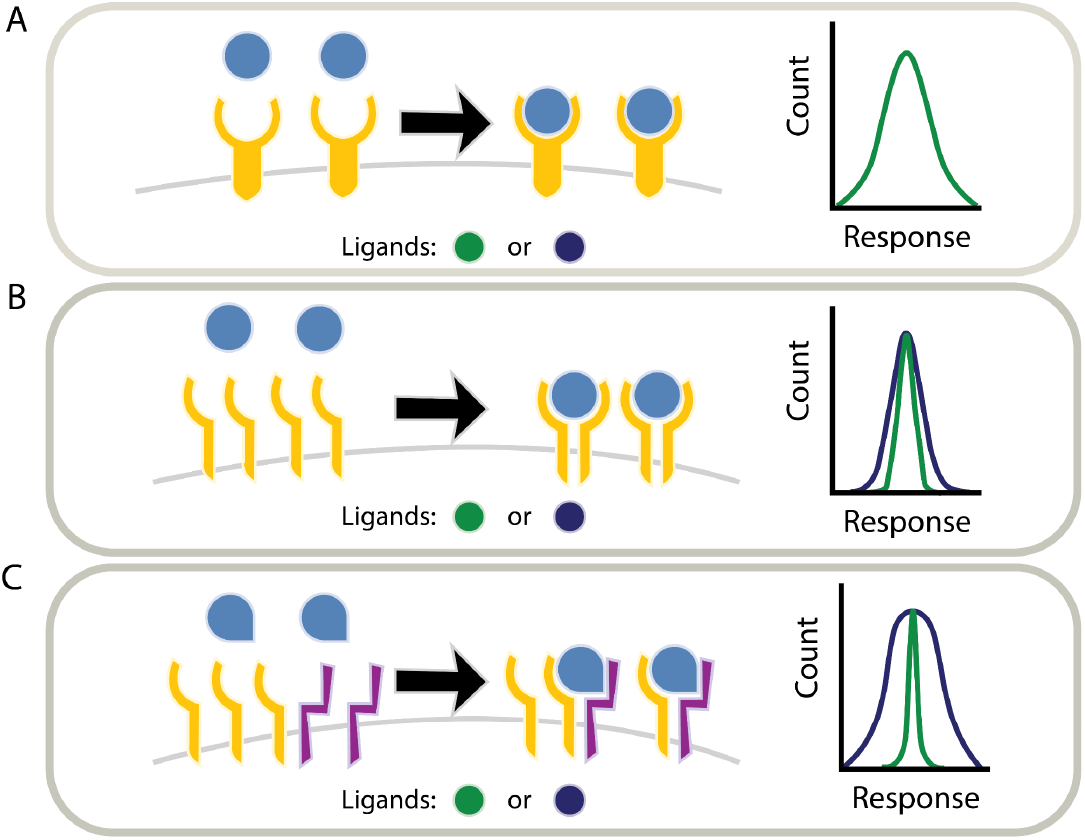
Different receptor architectures allow for different degrees of ligand discrimination based on response heterogeneity. (A) Receptors with a single-unit architecture always generate the same distribution of responses for a given mean response. Ligand identity can only affect the mean response but not the heterogeneity across the population. (B) The heterogeneity of the response for pathways containing homodimeric receptors can change by up to 2-fold, depending on the parameters of the ligands. In this way, ligands can be discriminated at the population level based on response heterogeneity. (C) Finally, pathways with heterodimeric receptors provide higher control over the response heterogeneity when the abundance of the two receptor subunits differ. This allows for a high degree of ligand discrimination for systems with an unbalanced expression of receptors.

These results provide a possible functional implication for multi-subunit receptor complexes that are often used in mammalian signaling pathways. A recent study [10] demonstrated that heterodimers might provide optimal ligand discrimination through distinct activation strength. Our work suggests a distinct possible functional advantage for such pathway interaction motifs, as they enable ligand-tunable variability. Under this hypothesis, when ligands are applied at a tissue level, such as interferon, TGFβ, BMP, or other cytokines, the responses are induced across the entire population of cells. In this case, controlling the heterogeneity of the response can provide a biological advantage, and the pathway would evolve to utilize multiple heterodimeric receptors. On the other hand, if the signals are inherently operating on individual cells, these pathways could utilize a single protein receptor. For example, in the juxtacrine NOTCH pathway, ligands are directed toward a single neighboring cell only. In agreement with our hypothesis, the Notch pathway has a single-unit receptor, and ligand discrimination in this pathway is achieved through ligand-distinct temporal dynamics [8]. Overall, we find that the choice of receptor type may depend on whether the pathway functions on a whole tissue or on individual cells locally.

The type I IFN pathway, which induces an innate immune response in tissues attacked by viruses [3], is a hallmark example of a multi-ligand pathway comprising, in humans, 16 ligand variants that are secreted to activate cell populations. Ligand discrimination by individual cells in the IFN pathway has been studied extensively [6,10,44]. However, many of the features of our model are exhibited by the IFN pathway suggesting a possible role for ligands in controlling the response heterogeneity. IFN receptor is composed of two subunits that promiscuously bind the 16 ligands [4,44]. Furthermore, previous studies have demonstrated that heterogeneity in response to the type I IFN is the result of heterogeneity in the receptor subunits’ expression levels throughout the cellular population, with the two subunits being expressed at distinct levels [14,29,45]. Moreover, different cell types expressing different ratios of subunits have been shown to have differences in heterogeneity in the response to the same type I IFN ligand in a dose-dependent manner [46]. These findings suggest that the IFN pathway is in a regime that gives rise to ligand-dependent heterogeneity. Indeed, recent studies have shown, in both the type I IFN pathway and other innate immunity-related cytokines [31,46], that heterogeneity in the response might have a functional role in controlling inflammation. As such, It would be interesting to test whether the spectrum of diverse type I IFN ligands results in distinct variability, adding to the control over the functional heterogeneity, which seems to play a major part in the immune response.

In this work, we concentrated on continuous graded responses to ligands. It has been demonstrated that in some pathways, the graded receptor activity induces a binary transcriptional response and that such a response is regulated at the population level [47]. In the same study, they further show that binary output can benefit from population heterogeneity [47]. For example, as we vary a signal-inducing heterogeneous complex formation, the percent of transcriptionally active cells within the population will gradually increase from zero to the entire population. For a homogeneous complex formation, the percent of responding cells will exhibit a switch-like behavior. Thus, ligand-dependent heterogeneity described in this paper could allow cells to control their activation mode.

Finally, cells in a population generally sense combinations of ligands within the same pathway interacting promiscuously with the receptors [5,15,16]. In particular, type I IFN is known to have 16-18 ligands that can all signal through the same set of receptors [6,44]. It would be interesting to extend our model to such cases of ligand combinations. In particular, using combinations of a high-variability inducing ligand with a low-variability inducing ligand might provide a way to regulate the heterogeneity of the response continuously.

Ligands mediate information between cells to generate robust, coordinated responses. Mostly, ligands are considered to act on individual cells and regulate their level of activity. We have shown that ligands can further determine population-level properties, such as the heterogeneity of the population independently of the average response level. Furthering our understanding of the capacity of ligands to regulate population-level features will bring us closer to understanding major biological processes such as the immune response, cancer, autoimmune diseases, and multicellular development. Moreover, a better understanding of controlling and manipulating heterogeneity within a population of cells would have implications in synthetic biology and artificial circuit design.

## Methods

We have developed analytical mathematical models for the different ligand-receptor architectures using binding-unbinding kinetic equations. Models were solved for the steady state response analytically, and the solutions were validated using the Equilibrium Toolkit (EQTK). EQTK is an optimized Python-based numerical solver for biochemical reaction systems [37,38]. Parameter scanning and simulations were performed using R version 3.6.3. Further details are provided in the supplementary text.

## Acknowledgment

Y.E.A is supported by the Israel Science Foundation (grant 1105/20) and by a research grant from the Sygnet Fund.

## Supplementary Information

### Receptor architecture mathematical models

We set out to develop mathematical models to describe the binding of ligands and the subsequent cellular responses for three different receptor architectures. We consider a basic model with a single unit receptor (the AL model), a model with a homodimeric receptor composed of two identical subunits (the ALA model), and a model with a heterodimeric receptor composed of two different subunits (the ALB model). Specifically, we focus on the level of ligand-receptor complex formation, leading to the activation of an intracellular signal mediator. The model does not contain other processes that might affect the cellular response, such as feedback loops, non-canonical signaling, and enzymatic signal amplification.

#### The AL model for ligand binding by a single-unit receptor architecture

In a single-unit receptor model, which we refer to as the AL model, we consider stimulation of the pathway by the ligand *L*, at a concentration *C*_*L*_. The ligand binds to a receptor, *A*, and forms a full complex *F*_*L*_. We consider a first-order kinetics where the forward binding rate *k*_*f L*_ and the reverse binding rate *k*_*r L*_ are intrinsic properties of the ligand and depend on its identity. This reaction can be summarized as

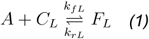

In addition, for all models we assume that the volume for the ligands is large so there are significantly more ligand molecules than receptors, or *V* → ∞. Under this assumption, ligand concentrations remain constant, such that the initial concentration, *C*^*0*^_*L*_, doesn’t change and

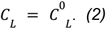

With this we can write the dynamical equations that describe the ligand binding by a single unit receptor:

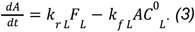

Here *C*^*0*^_*L*_ denotes the concentration of the ligand in volume *V*, while *A* and *F*_*L*_ denote the absolute number of receptors and complexes on the surface of a cell. We assume that production and consumption of the different molecules are in steady state, allowing us to neglect the endocytosis of ligands and receptors. The conservation of mass requires that the total number of each type of molecule remains constant, regardless of whether it is free or in a complex with other species. Denoting the initial receptor level as *A*^*0*^ we obtain

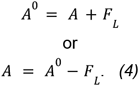

As changes in the ligand binding by receptors occur much faster than changes in receptor expression [5], we consider the behavior of the system to be at steady state. As such, all time derivatives become zero, and equation *3* can be solved to give:

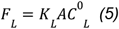

where *K*_*L*_ is defined as 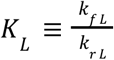, and describes the affinity of ligand *L* to receptor *A*. Plugging equation *4* into *5* we can solve the behavior of complex *F*_*L*_ in steady state

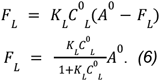

This equation provides the dependence of the number of full complexes on the three model parameters, and simulating the parameters gives the expected Michaelis-Menten relationship (supplementary Figure 1A) [6,10].

#### The ALA model for ligand binding by a homodimer receptor

We next consider an architecture with two identical receptor subunits that homodimerize upon ligand binding, which we refer to as the ALA model. We consider a ligand *L*, at a concentration of *C*_*L*_ that binds to a receptor subunit, denoted by *A*, and forming a partial complex, *P*_*L*_. This partial complex further binds a second subunit to form the full complex *F*_*L*_. As before, we assume that the binding is reversible with first-order kinetics. We consider the forward and reverse binding rates for the formation of the partial complexes, *k*^*P*^_*f L*_ and *k*^*P*^_*r L*_, and full complexes *k*^*F*^_*f L*_ and *k*^*F*^_*r L*_, to be intrinsic properties of the specific ligand variant. These reactions can be summarized as

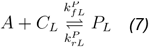

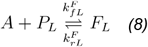

We use *C*^*0*^_*L*_ to denote the concentration of the ligand while *A, P*_*L*_ and *F*_*L*_ denote the number of receptors and complexes on the surface of a cell. As above, we consider the concentration of the ligand to remain constant throughout the reaction (equation 2). We can now write the dynamical equations resulting from these reactions (7,8):

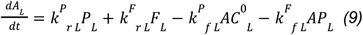

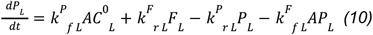

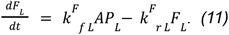

Considering conservation of mass, and denoting the initial values of the receptor subunits as A^0^, we obtain

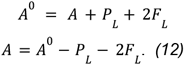

Note that since a single complex, *F*_*L*_, comprises two receptor units it appears in the conservation of mass equation with a factor of two.

The steady state solution for this model is achieved by equating equations *9* and*10* to zero. Solving the resulting system of equations we find

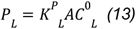

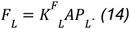

Where *K*^*P*^_*L*_ is defined as 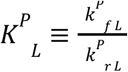, and *K*^*F*^_*L*_ is defined as 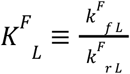, denoting the affinity of the ligand and ligand-bound receptor subunit to subunit *A*. By plugging equation *12* into equations *13* and *14*, we can obtain the dependence of *P* on the parameters in steady state:

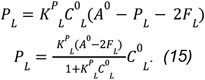

Finally, by plugging this into equation *14* we can arrive at a quadratic equation for *F*_*L*_,

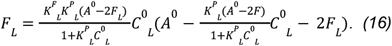

This can be solved to find

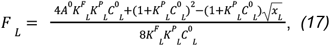

where *x*_*L*_ is defined to be

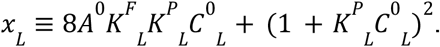

Equation *17* can be solved numerically for any given set of parameters, and doing this gives the expected non-monotonic relationship (supplementary Figure 2A) [35,36].

We note that the quadratic equation *16* has two solutions with a negative or positive sign for the square root in equation *17*. However, taking the positive solution results in values for F_L_ that are higher than *A/2* and thus this solution is not biologically relevant. We therefore focus on the negative solution for the square root.

#### The ALB model for ligand binding by a heterodimer receptor

A third model we consider in the paper is based on an architecture with two different types of receptor subunits that heterodimerize upon ligand binding.

We consider a sequential complex formation with a ligand, *L*, at a concentration *C*_*L*_, that binds first to a specific receptor type, *A*, to form a partial complex, *P*_*L*_. As a second step, the partial complex further binds a second receptor subunit, *B*, to form the full complex *F*_*L*_. As with the previous models, we assume that the binding is reversible with first-order kinetics. The forward and reverse binding rates for the formation of the partial complexes, k^P^_f L_ and k^P^ _rL_, and full complexes k^F^_f L_ and, k^F^ _rL_, are considered as intrinsic properties of the specific ligand variant. Using these notations, the reactions can be summarized as (cf equations *7*,*8*):

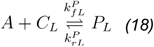

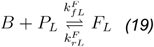

We further denote, as before, the initial ligand concentration by *C*^*0*^_*L*_, which is considered to remain constant. *A, B, P*_*L*_ and *F*_*L*_ denote the number of receptors and complexes on the surface of a cell. We can thus write the dynamical equations describing these reactions as:

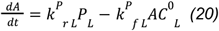

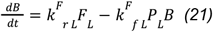

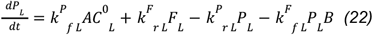

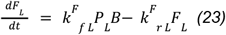

As discussed previously, considering conservation of mass, and denoting the initial values of the receptors *A*^*0*^ and *B*^*0*^ we obtain

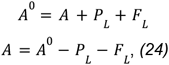

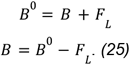

We consider our system in steady state, such that all time derivatives vanish, and the system can be reduced to two equations, which can be solved to give

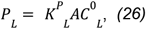

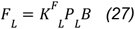

where *K*^*P*^_*L*_ and *K*^*F*^_*L*_ are defined as 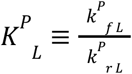 and 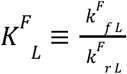, describing the affinity of the ligand to subunit *A*, and the affinity of the partial complex to subunit *B*, respectively. Plugging equation *24* into equations *26*, we obtain the *P*_*L*_ dependence on the parameters in steady state:

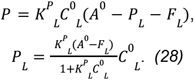

This can be plugged into equation *27* together with equation *25* and give rise to a quadratic equation for *F*_*L*_:

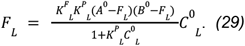

Solving equation *29* we get

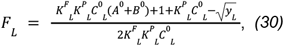

where *y*_*L*_ is defined to be:

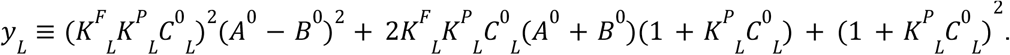

Equation *30* can, like in the case of the previous model, be solved numerically for any given set of parameters.

As above, we chose a specific sign for the square root in equation 30 to provide the biologically relevant solution where *F*_*L*_ does not increase beyond the level of *A*^*0*^ or *B*^*0*^.

#### Calculating the intracellular response

The models enable us to determine the amount of full complexes that are generated given a specific ligand, with given parameters. However, in many signaling pathways the binding of a ligand to its receptor is only the first in a series of steps that culminate in the cell’s response to the ligand, usually in the form of changes in gene expression [3,5,8]. We consider a simplified model for the downstream signal transduction where the full complex, *F*_*L*_, elicits gene expression at a specific rate, *e*_*L*_. This parameter can depend on the identity of the complex and in particular, in our model it is determined by the ligand. The *on* and *off* rates for a specific ligand as well as the physical proximity of the two receptor subunits resulting from the ligand dependent interaction could affect *e*_*L*_ independently of other ligand dependent parameters.

Given *e*_*L*_, we can calculate a cell’s response, *E*_*L*_, as a function of the amount of full complexes formed assuming linear dependence:

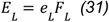

By plugging the solutions for equations *6, 17* and *30* into equation *31*, we can solve for the ligand dependent cell response of the different receptor architecture models.

## Local Scaling

### Calculating local scaling

In order to quantify the effect of ligand parameters on the distribution of responses one needs to assume a specific distribution of receptors in the population and perform full simulations on the entire population. However, we aimed to define a local measure that would enable us to calculate a metric for a single cellular configuration that would provide a handle on the global diversity in the response. To achieve this, we defined a basic metric of local scaling (*S*) describing the local power law dependence of the response on the receptor amount:

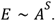

For infinitesimal changes, the local scaling can be calculated as the derivative of log(*E*) as a function of log(A):

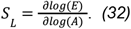

Intuitively, this metric measures the variability in ligand induced cell responses, *E* as we vary specific parameters, in particular the receptor levels on the cell membrane. We further wanted this metric to reflect effects of relative changes in receptor levels on relative changes in the response.

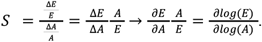

As the exponent in the power law increase, the same variability in the distribution of receptors will be translated into a higher dependence in the response of the system (Figure S1B). Thus, the local scaling metric can be used to estimate effects on the global variability at the population level.

### Calculating scaling with respect to receptor amounts in the models

We first consider the AL model’s response, given from plugging equation *6* into equation *31*

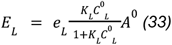

We can next break the calculation of the response’s scaling with respect to *A* into two parts, first calculating the derivative of 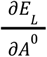, then its multiplication by 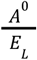. Solving the derivative of 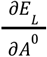 we obtain

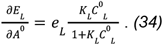

Next we multiply the derivative by 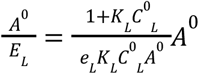, and obtain the scaling

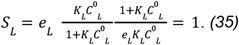

Thus, for the single unit receptor, the scaling is constant and independent of ligand parameters (see main text and Figure 2 for details).

In the same fashion we can calculate the scaling of the response of the ALA model with respect to changes in the initial subunit receptors amount. First, calculating the derivative of 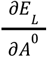 gives

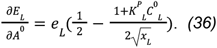

Next we multiply by 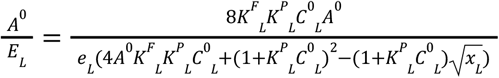, as before, and obtain

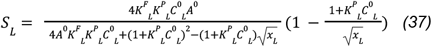

Which can be numerically simulated for any given set of parameters. The *x*_*L*_ in equations *36* and *37* is defined in equation *17*.

Finally we calculated the scaling of the response in the ALB model with respect to changes in the total amount of the receptor subunits. In this case, however, as there are two different types of receptor subunits, we need to calculate the scaling with respect to each receptor separately. Starting with receptor subunit *A*, we can follow the same process as for the previous model, for which we obtain

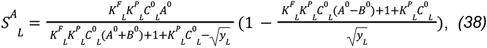

and following the same process for subunit *B*, we obtain

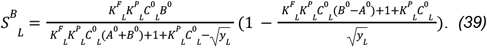

Both can be numerically simulated for any given set of parameters. The *y*_*L*_ in both equations *38* and *39* is defined in equation *30*.

Importantly, we note that, based on the calculations, the scaling of the response to changes in the receptors or receptor subunits is independent of the activity rate, *e*_*L*_, in all models.

## Reducing parameter space using nondimensionalization

Our models include several parameters that determine the response of the system. For example, in the basic model for a single unit receptor we have four parameters, including three ligand parameters: concentration (*C*_*L*_), affinity to the receptor (*K*_*L*_) and activation rate (*e*_*L*_), as well as a single cellular parameter, the amount of receptors (*A*). In order to fully determine the behavior of a model across all parameter values one needs to analyze the solution across the entire parameter space. Nondimensionalization is a standard method to reduce the number of parameters that should be independently varied in order to fully analyze the system [5]. When studying the biochemical parameters we should notice that they are dimensional, however the units can be chosen arbitrarily. This choice of units does not affect the normalized behavior of the system. For example changing the units of concentration from mg/ml to ng/ml would change the numeric value of *C*_*L*_ and *K*_*L*_ but will not change the resulting number of complexes.

More generally, in the AL model, *C*_*L*_ has units of ligand concentration and *K*_*L*_ has units of inverse ligand concentration. Changing these units by a scaling factor *α* will affect the values of concentrations and binding affinities in the following way:

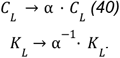

However, such coordinated change will result in exactly the same values for the number of full complexes, *F*_*L*_. While the value of each parameter is changed in the new unit scheme, the product *K*_*L*_*C*_*L*_ is dimensionless and will not change under redefinition of the units. From the equation for *F*_*L*_ (equation *6*) we see that the parameters only appear in this specific combination. Therefore, it is enough to only consider the case *K*_*L*_ = 1 (or alternatively *C*_*L*_ = 1) as other values can be mapped to this one by the transformation in equation *40*. Intuitively, we can always define the units of measurements for the ligand concentration such that the EC50 concentration is one. In these units *K*_*L*_ is equal to one, and the only relevant parameter is the ligand concentration. This process of nondimensionalization dictates that only these parameters will affect the behavior of the system. When scanning the parameter space we will thus only analyze such dimensionless parameter combinations. Overall for the AL model, there are three relevant parameters: *K*_*L*_*C*_*L*_, *e*_*L*_ and *A*.

Doing the same procedure for the ALA model, we start with four ligand parameters: *C*_*L*_, *K*^*P*^_*L*_, *K*^*F*^_*L*_ and *e*, as well as one cellular parameter: *A*. In this case we consider the dimensionless combination *C*_*L*_*K*^*P*^_*L*_, as before. In addition, by considering a similar change in the units of receptor amount, we get another dimensionless combination, *K*^*F*^_*L*_ *A*. Overall the ALA model has three relevant parameters: *K*^*P*^_*L*_ *C*_*L*_, *K*^*F*^_*L*_ *A* and *e*_*L*_.

Finally, for the ALB model, we start with four ligand parameters: *C*_*L*_, *K*^*P*^_*L*_, *K*^*F*^_*L*_ and *e*_*L*_, and two cellular parameters: *A* and *B*. Using the same unit transformation for ligands and receptors, we will consider the following dimensionless parameters: *K*^*P*^_*L*_*C*_*L*_, *K*^*F*^_*L*_*(A+B), A/B* and *e*_*L*_.

## Relationship between F_L_ and S_L_ in the ALA model

To determine the functional relationship between the full complex and the scaling in the ALA model we first define the following non dimensional parameters:

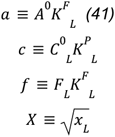

Using these parameters,equation *17* can be rewritten as:

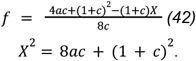

Similarly, *S*_*L*_ (equation *37*) becomes:

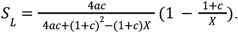

We consider the inverse of *S*_*L*_:

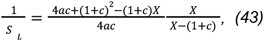

and, with some algebra, this can be rewritten as:

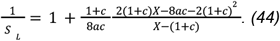

Further development of equation *44* gives us:

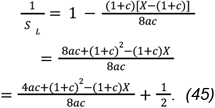

Using equation 42, we find

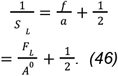

Inverting this equation we find a simple dependence between *S*_*L*_ and *F*_*L*_ (Figure 3C):

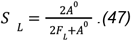

## The scaling of the full complex across the parameter space of the ALA model

The metric defined in equation *32* is defined as local scaling of the number of full complexes, *F*_*L*_, with the initial amount of receptor subunits, *A*^*0*^. Here we will study the equation for *F*_*L*_ (equations *17* or *42*) to determine this scaling and its dependence on *K*^*F*^_*L*_*A*^*0*^. We first consider the two regimes discussed in the main text, in both of which *c* = 1, such that the ligand is supplied at its EC50 (equation *44*). Under this condition we obtain:

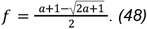

We study two regimes: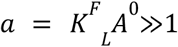 and 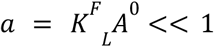, as described in the main text. For the first regime, *a*≫1, equation *48* can be simplified to be

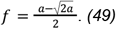

As 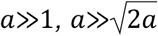. Thus, equation *52* can be further simplified to

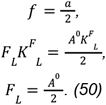

Thus, when *K*^*F*^ *A*^*0*^≫1 the full complex *F*_L_ scales linearly with the initial amount of the receptor subunit *A*^*0*^.

In the second regime, *a* << 1, we can expand equation *49* by using the taylor expansion for a square root. We find:

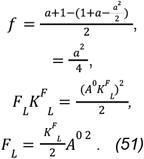

Thus, when *K*^*F*^_*L*_ *A*^*0*^<<1, *F*_*L*_ scales quadratically with the number of receptors, A^0^.

We can further extend our findings to a general ligand concentration. Here, under the first condition, when 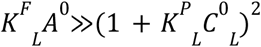, we obtain:

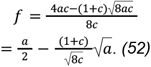

In this regime, the equation can be approximated as

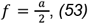

so that

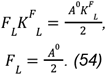

Thus, we find that even in this more general case *F*_*L*_ scales linearly with *A*^*0*^.

Next, when looking into the the second regime where *K*^*F*^_*L*_*A*^*0*^*<<1*, and expanding equation *42*, we obtain:

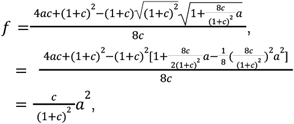

Which can be rewritten as

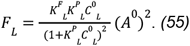

Thus, under the condition of *K*_*L*_^*F*^ *A*^*0*^*<<1*, regardless of the *K*^*P*^_*L*_ *C* ^*0*^_*L*_, *F*_*L*_ scales quadratically with the number of receptors, A^0^.

## Unordered ligand binding in the ALB model

When modeling the ALB, heterodimeric receptor architecture, we assume that the assembly of the full complex is sequential. In this case, the ligand (*L*) first binds a specific receptor subunit type (*A*) and only then the partial complex binds the second receptor subunit type (*B*). While this is described as the mode of operation of pathways such as type I IFN, TGFβ, and BMP *in-vivo [4*,*5*,*39]*, under certain conditions the binding might proceed in parallel. Therefore, we consider the case of unordered binding in the ALB model. The resulting reactions can be summarized as:

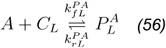

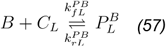

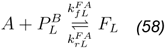

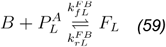

with *C*_*L*_ being the ligand concentration as before, and *P*^*A*^_*L*_ and *P*^*B*^_*L*_ being the partial complexes generated through *L* binding either *A* or *B* respectively. The new binding reactions are considered to have first-order kinetics. Here, *k*^*PA*^_*f L*_ and *k*^*PA*^_*r L*_ are the forward and reverse binding rates of the ligand to receptor subunit *A* respectively, and similarly *k*^*PB*^_*f L*_ and *k*^*PB*^_*r L*_ are the forward and reverse binding rates of the ligand to receptor subunit *B*. Likewise, *k*^*FA*^_*f L*_ and *k*^*FA*^_*r L*_ are the forward and reverse binding rates, respectively, of *P*^*B*^ and subunit *A*, while *k*^*FB*^_*f L*_ and *k*^*FB*^_*r L*_ are the forward and reverse binding rates, respectively, of *P*^*A*^ and subunit *B*. These parameters are all intrinsic properties of the ligand.

As before, assuming steady state and using equation *2*, we find the following three equations:

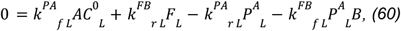

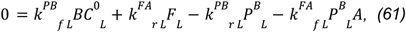

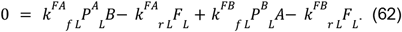

As there is no energy invested in the formation of these complexes, e.g, in the form of ATP hydrolysis, this indicates that the kinetic rates satisfy detailed balance relation. Defining the affinities as

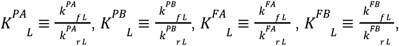

detailed balance can be written as

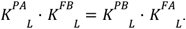

With this we obtain:

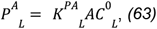

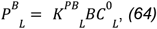

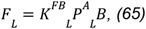

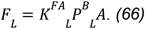

The condition of detailed balance also dictates that one of the above equations is redundant. As such, we can remove equation *65*.

Next, considering conservation of mass, and denoting the initial values of the receptors *A*^*0*^ and *B*^*0*^ we obtain

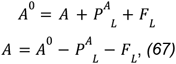

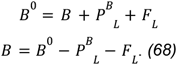

By plugging equations *67* and *68* into equations *63, 64* and *66*, we can obtain the dependence of the two partial complexes’ behavior on the parameters in steady state:

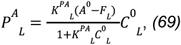

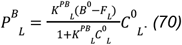

Finally, by plugging these into equation *66* we can arrive at a quadratic equation for *F*_*L*_,

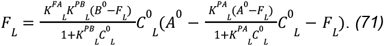

This can be solved to find

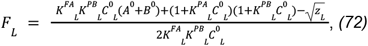

where *z*_*L*_ is defined to be:

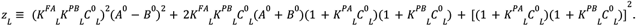

Equation *72* can, like in the case of the previous models, be solved numerically for any given set of parameters.

Next, using the same process described above, we can find the scaling for this model with respect to changes in the initial amount of the receptor subunits. First to subunit *A*:

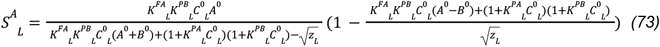

And following the same process for subunit *B*, we obtain

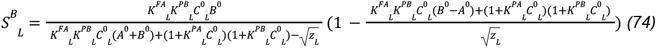

Solving these equations for different parameters, we can show that the order in which the ligand binds the receptor subunit changes neither the range of the different scalings (supplementary Figure 3G), nor the molecular mechanism behind the diversity in scaling values.

**Supplementary figure 1.**
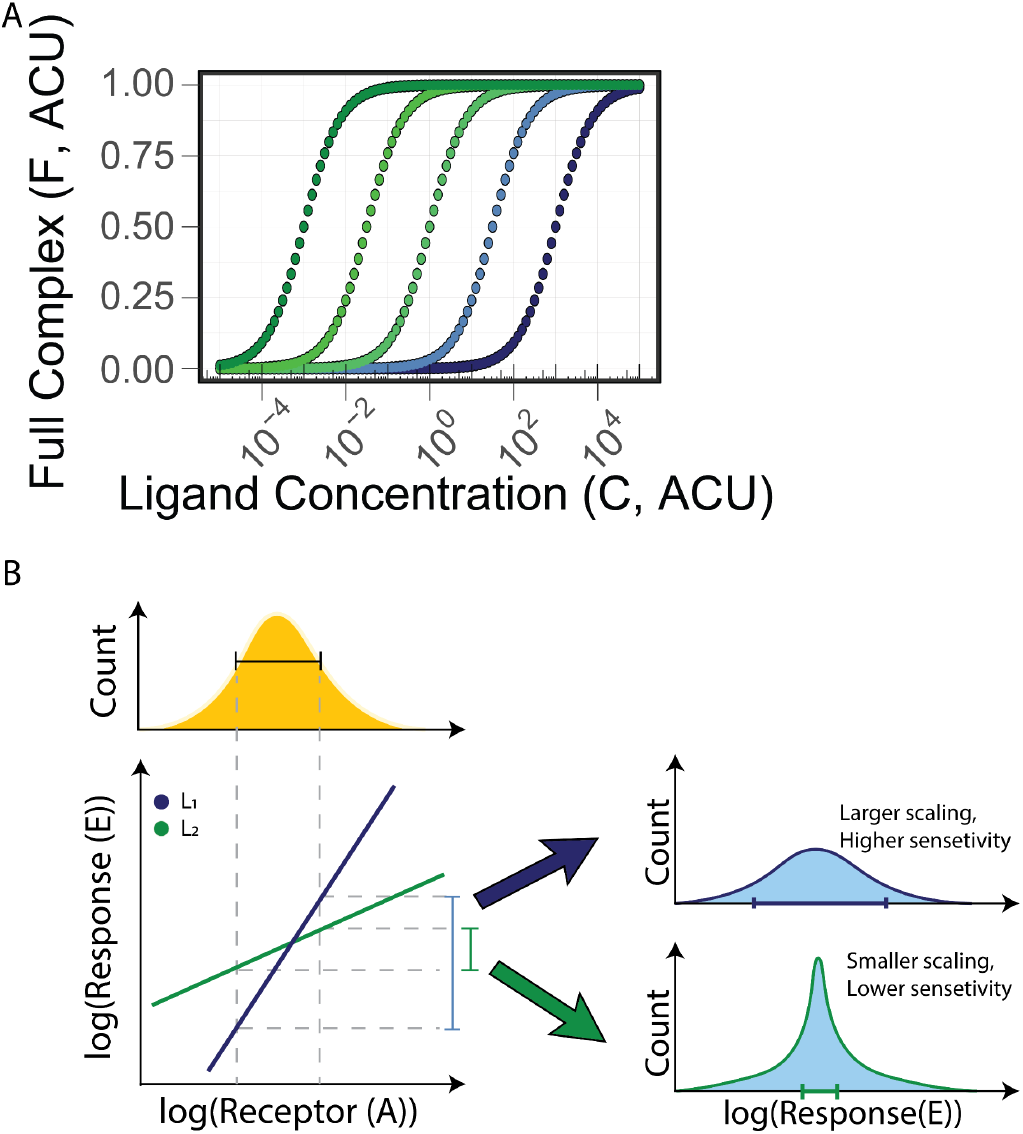
Single-unit receptor response and scaling. (A) We consider The formation of *F*_*L*_ for the single unit receptor pathway given the five different ligands described in Figure 1B over multiple concentrations and *A*^*0*^ *= 1*. (B) The dependence of the response on the receptors is plotted for two systems with either high (blue) or low (green) scaling. Given a specific distribution of receptors in a population of cells, the high-sensitivity system (blue) with larger scaling will generate a highly variable response. In contrast, the low-sensitivity system (green) has lower scaling and will generate responses with lower variability. ARU = Arbitrary Receptor Units, ACU = Arbitrary Concentration Units.

**Supplementary figure 2.**
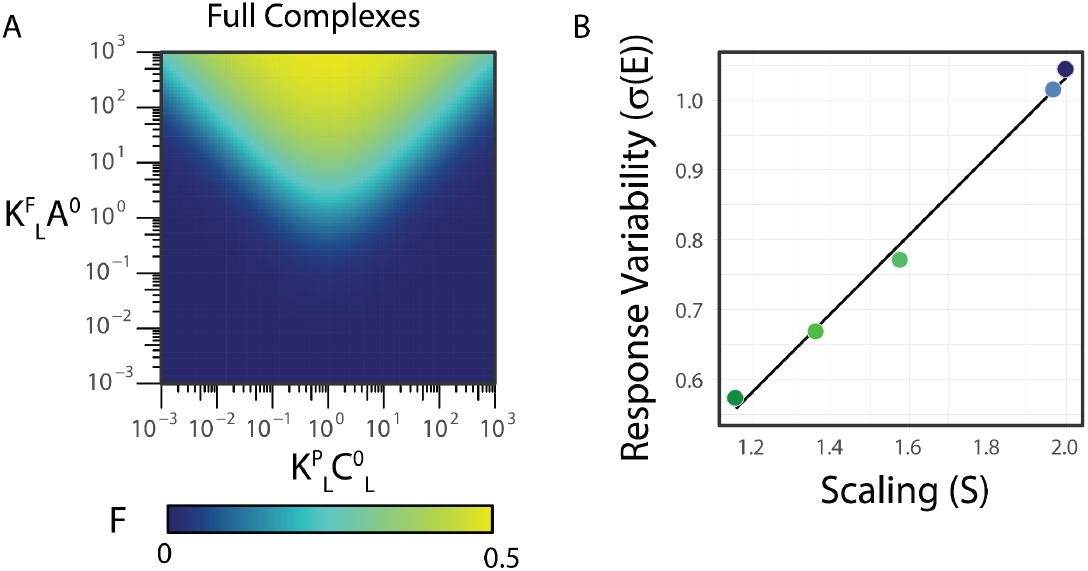
Homodimeric receptor response heterogeneity correlates with its scaling to subunit abundance. (A) The homodimeric model’s full complex (*F*_*L*_) is plotted across model parameters showing a non-monotonic response to ligand concentrations. (B) The standard deviation in the response was calculated (cf. figure 4C) and plotted for ligands with different scaling values. Different ligands are colored as in Figure 4B. The relationship can be approximated by a linear dependence (rho = 0.997, p = 0.0001683, Pearson correlation).

**Supplementary figure 3.**
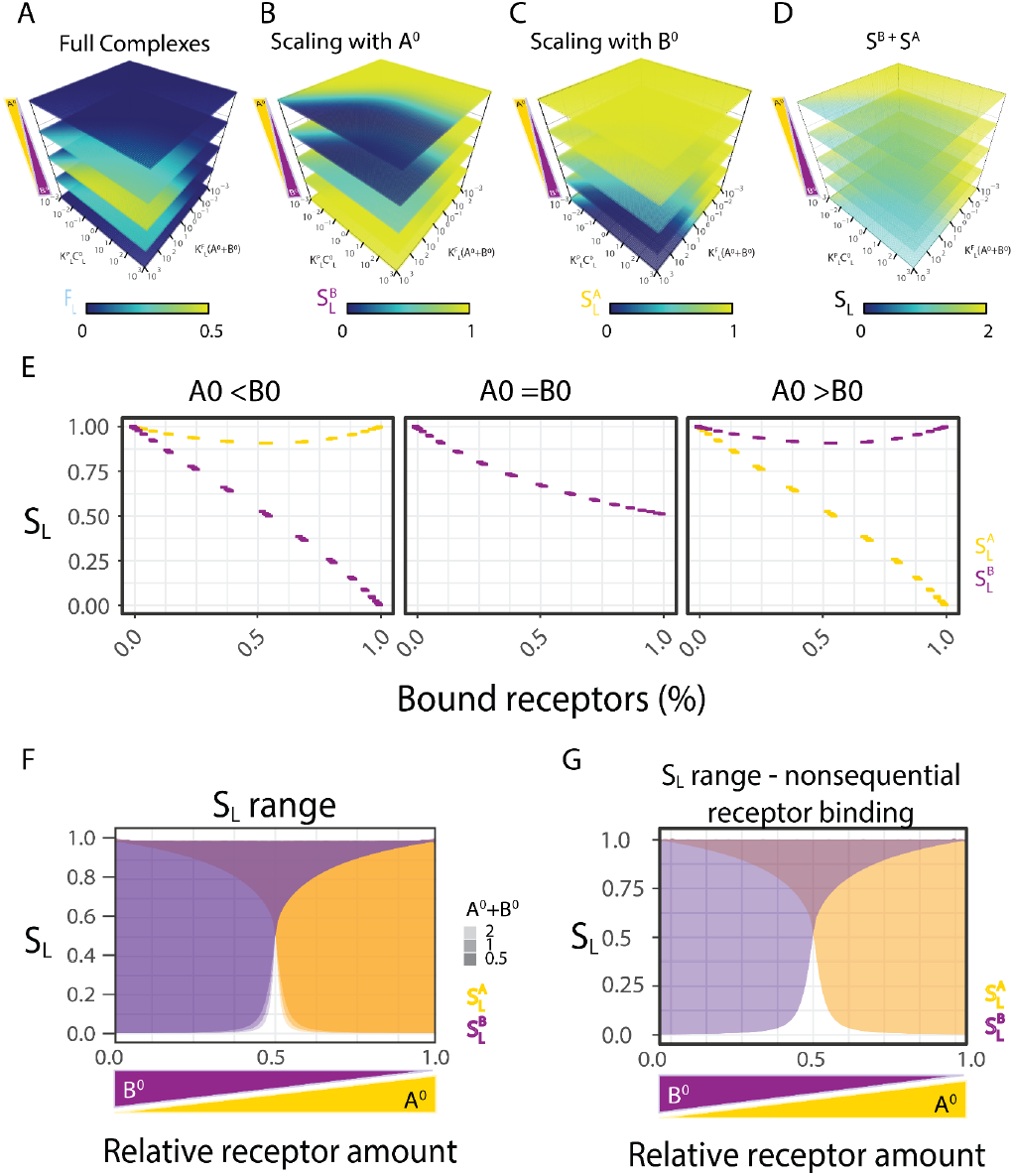
Heterodimeric receptor architecture extends the ligand-dependent control of response heterogeneity. (A - C) The sequential binding heterodimeric receptor model’s full complex (*F*_*L*_) (A) and the scaling with receptor subunits *A*^*0*^ and *B*^*0*^ (B and C, respectively) are shown across the model’s dimensionless parameters as discussed in Figure 5B,C and across five different ratios of the receptor subunits *A*^*0*^/*B*^*0*^ [0.001, 0.25, 0.5, 0.75, 999]. (D) Total scaling is determined by the addition of the scaling with *A*^*0*^ and *B*^*0*^ as shown in B and C. (E) The scaling with each subunit is determined by its given relative amount and the fraction of bound receptors (*F*_*L*_/[the less abundant subunit]). This dependence is plotted for different ratios of subunits *A*^*0*^ and *B*^*0*^. (F) Range of the model’s scaling with the receptor subunits (*S*^*A*^_*L*_ in orange and *S*^*B*^_*L*_ in purple), given different total amounts of the receptor subunits (shades of orange and purple). The range was calculated under the same model parameters and subunit ratios as in figure 5D. (G) Range of the heterodimeric receptor with non-sequential subunit binding model’s scaling with the receptor subunits (*S*^*A*^_*L*_ in orange and *S*^*B*^_*L*_ in purple) under different model parameters as dependent on the relative amount of the receptor subunits from 0.001 to 999. Given a range of dimensionless parameters, the resulting scaling *S* retains an extensive range in the same manner as the sequential model.

**Supplementary Figure 4.**
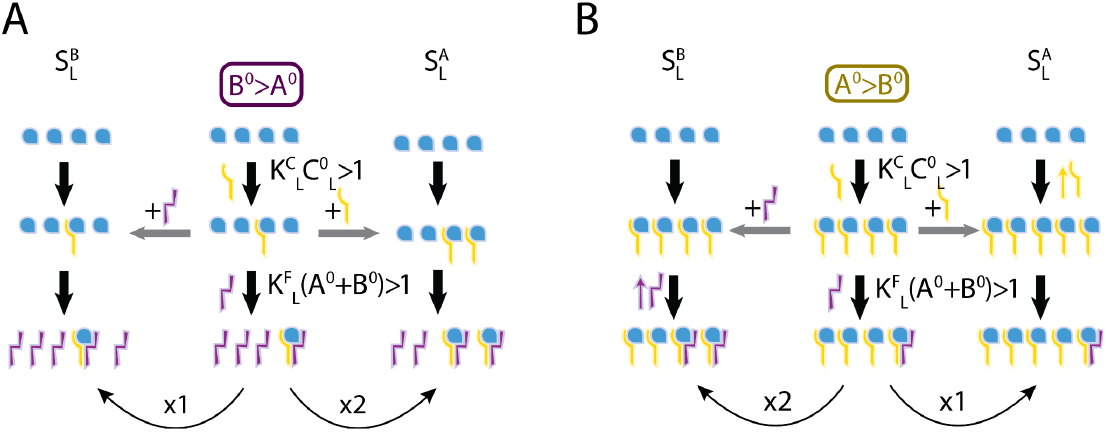
Cartoon representing the molecular mechanism behind the dependence of the scaling on the relative receptor amounts in the high-affinity regime. (A) When *B*^*0*^ is larger than *A*^*0*,^ and affinities are high (ligand concentrations are saturating, red circles in Figure 5B), all free ligands will bind directly to *A* (yellow), leaving no free subunit, and all *B* will bind to any free *P*_*L*_. As *A*^*0*^>*B*^*0*^, there are more free *B* than free *P*_*L*_, making the complex amount *F*_*L*_ insensitive to *B* and dependent on *A*. (B) Alternatively, when *A*^*0*^ is more abundant than *B*^*0*^, all B subunits will form a full complex, *F*_*L*_. In this case, *F*_*L*_ is strongly dependent on *B* and insensitive to *P*_*L*_ and, thus, to *A*.

## Notes

### Competing Interest Statement

The authors have declared no competing interest.

